# Neural network approach to somatic SNP calling in WGS samples without a matched control

**DOI:** 10.1101/2022.04.14.488223

**Authors:** Sergey Vilov, Matthias Heinig

## Abstract

Somatic variants are usually called by analysing the DNA sequences of a tumor sample in conjunction with a matched normal. However, a matched normal is not always available for instance in diagnostic settings. To unlock such data for basic research single-sample somatic variant calling is required. Previous approaches can not easily be applied in the case of typical whole genome sequencing (WGS) samples.We present a neural network-based approach for calling somatic single nucleotide polymorphism (SNP) variants in tumor WGS samples without a matched normal. The method does not require any manual tuning of filtering parameters and can be applied under the conditions of a typical WGS experiment. We demonstrate the effectiveness of the proposed approach by reporting its performance on 5 SNP datasets corresponding to 5 different cancer types.

The proposed method is implemented in Python 3.6 and available as a GitHub repository at https://github.com/heiniglab/deepSNP.

## 1 Introduction

Somatic variants are genetic alterations accumulating in non-germline cells during the lifetime of the organism. Somatic mutations that corrupt genes regulating cell growth, programmed cell death or neovascularization can promote the formation of a cancer (Li *et al*., 1992; Hanahan and Weinberg, 2000; Young *et al*., 2013; Gao *et al*., 2014). Hence, identifying somatic mutations is important to understand cancer genesis, choose treatment strategies and make prognosis.

The broad spectrum of somatic variants comprises single nucleotide polymorphism (SNP) variants, short insertion and deletion (INDELs), large copy number alterations, and structural rearrangements (Ciriello *et al*., 2013). All these mutations can be detected via somatic variant calling.

Somatic variant calling should accomplish two tasks. First, true variants should be separated from sequencing and alignment artefacts. Second, somatic variants should be distinguished from germline mutations, which should be identified and filtered out. In popular somatic variant calling pipelines, such as GATK (McKenna *et al*., 2010) and Strelka (Saunders *et al*., 2012), these tasks are accomplished by constructing alignments of tumor and normal reads and removing artefacts by means of statistical models and filtering rules tailored to remove specific artefacts. In recent years, it has been shown that better results can be achieved using machine learning techniques.

For example, the method called Cerebro (Wood *et al*., 2018) is based on extremely randomized trees trained on two sets of features derived from tumor-normal alignment with two different alignment programs. Cerebro has been reported to detect somatic variants with a better accuracy compared to conventional callers. Deep learning-based methods, such as NeuSomatic (Sahraeian *et al*., 2019) and DeepSSV (Meng *et al*., 2021), have also demonstrated encouraging results.

However, a matched normal is not always available. This is a very common scenario in retrospective analysis of samples from clinical trials, pathology archives, and legacy biobanks. Absence of a patient’s consent or financial restrictions can also prevent collection of the normal sample. Removing artefacts when calling somatic variants without a matched normal is similar to removing artefacts when calling germline variants. First, reads with low quality scores are to be removed. Afterwards, germline variants and artefacts can be separated based on their variant allele fraction (VAF): germline variants are expected to have VAF around 50%(heterozygous) or 100%(homozygous) whereas artefacts are usually associated with much lower VAF values. Although statistical models have long been used for artefact filtering in germline variant calling pipelines (Xu, 2018), some recent studies have argued that machine learning techniques can provide a superior filtering quality without the need to tailor specific filtering rules.

For example, in the pioneering Deep Variant approach (Poplin *et al*., 2018), candidate variants in the form of piled up read images were presented to a convolutional neural network. The neural network output was then used to judge whether the input alteration was a true germline variant or an artefact. For true variants, zygosity was also predicted. Deep Variant was reported to outperform filtering tools proposed in conventional germline variant calling pipelines, such as GATK (McKenna *et al*., 2010) and Strelka (Saunders *et al*., 2012). Later on, an even better performance was achieved for an alternative neural network architecture and a slightly different variant encoding scheme (Friedman *et al*., 2020).

In contrast to germline mutations, somatic variants often have a very low VAF due to tumor-normal contamination or tumor heterogeneity (Xu, 2018). This makes it difficult to separate somatic mutations from artefacts. Therefore, highly accurate statistical modeling and advanced error correction techniques, such as those based on machine learning, would be extremely valuable.

Separating somatic and germline variants when a matched normal is not available is also challenging. Since candidate variants can not be tested against a control sample, one can judge about a given variant only by comparing its characteristics with *a priori* information about somatic and germline mutations.

For instance, germline variants can be filtered out based on their VAF (Li *et al*., 2017): as mentioned above, VAF of somatic variants is often lower compared to their germline counterparts. This is the core idea of the method called SomVarIUS (Smith *et al*., 2016). To separate somatic and germline variants, it builds a beta-binomial model of the germline VAF distribution. Artefacts are filtered out using another statistical model which uses read base quality scores. The error in VAF estimation decreases with the read depth, so SomVarIUS needs a relatively high coverage (≥100x is recommended) to call somatic variants accurately enough. Such coverage is not typical in routine whole genome sequencing (WGS) experiments (Björn *et al*., 2018). It should also be noted that SomVarIUS relies on manually fixed filtering parameters.

Information about the mutational context can also be useful for germline variant filtering in single-sample somatic variant calling. For SNP variants, the mutational context is represented by the alternative allele and a sequence region of interest (ROI) around the variant position in the reference genome. Among other features, the mutational context is used by the machine learning-based approach called ISOWN (Kalatskaya *et al*., 2017) that can be employed to separate somatic and germline variants. To perform variant classification, ISOWN also considers information from public variant databases, variant effect prediction, sample frequency, VAF, etc. It is worth mentioning that ISOWN is unable to filter out artefacts. In addition, ISOWN has not been validated on WGS variants that might be absent in public variant databases and whose effect might be challenging to predict.

In this study, we present a new approach to identifying somatic SNPs in tumor WGS samples without a matched normal. The proposed method relies on recent advances in artefact filtering as well as on state-of-the-art approaches to germline variant removal in single-sample calling. In the core of the method is a neural network classifier trained using 3D tensors consisting of piledup variant reads. Trained on a number of WGS samples for which somatic and germline SNPs are provided as labels, the proposed method can then be used to identify somatic variants in previously unseen samples of the same underlying disease. The proposed method does not require manual parameter tuning and can be effective when applied to samples with a typical WGS coverage of around 30x.

In this paper, we evaluate the proposed method on 5 different cancers datasets. The resulting performance is then interpreted by analysing mutational signatures and VAF distributions of true SNP variants and artefacts. After that, we review our choice of the neural network architecture and discuss the impact of the dataset size as well as the sequence ROI length around the variant site.

## 2 Materials and Methods

### 2.1 Input data and preprocessing

The neural network classifier has been trained and evaluated on each of 5 datasets representing 5 different cancers: TCGA-LAML (acute myeloid leukemia), BLCA-US (bladder urothelial cancer), ESAD-UK (esophageal adenocarcinoma), LINC-JP (liver cancer), GACA-CN (gastric cancer). Four of these datasets (BLCA-US, ESAD-UK, LINC-CP, GACA-CN) were downloaded from the ICGC portal (https://dcc.icgc.org), the TCGA-LAML dataset being downloaded from the TCGA portal (https://portal.gdc.cancer.gov). Train/evaluation data for each dataset was generated based on tumor WGS samples. The total number of tumor samples used, the number of somatic SNPs per sample, the percentage of somatic SNPs in gnomAD v. 3.1.2 database (Karczewski *et al*., 2020), and the average tumor coverage are shown in Table 1 for each dataset. Mutational signatures of somatic variants are shown in Figure S1 for each dataset. Figure S2 illustrates mutational signatures of artefacts and germline variants. Variant allele fraction (VAF) distributions in each dataset are demonstrated in Figure S3.

**Table 1.** Description of the datasets considered in this study.

| Dataset | Tumor WGS samples | Somatic SNPs per sample | Somatic SNPs in gnomAD v. 3.1.2 | Average coverage |
| --- | --- | --- | --- | --- |
| TCGA-LAML | 47 | 393 | 30% | 35x |
| GACA-CN | 25 | 10230 | 18% | 37x |
| LINC-JP | 28 | 8529 | 13% | 54x |
| ESAD-UK | 41 | 53212 | 13% | 67x |
| BLCA-US | 23 | 17998 | 14% | 36x |

For supervised learning of a neural network classifier, labelled exemplars of both positive (somatic variants) and negative (germline variants plus artefacts) classes should be provided. To obtain positive class instances, a VCF file with pre-called somatic SNPs was downloaded for each WGS sample from the corresponding resource. For ICGC samples, we considered consensus VCF files from the ICGC portal. From each consensus VCF file we selected only variants on which all callers agreed. For the TCGA-LAML dataset, we considered somatic variants reported in (Cancer Genome Atlas Research Network, 2013). To obtain negative class instances, we ran the Mutect2 caller (Benjamin *et al*., 2019) (bundled with GATK v4) for each WGS tumor sample using default settings and without a matched normal. Known somatic variants were then removed from the Mutect2 output. Although identifying germline variants in the Mutect2 output was not necessary to train the neural network classifier, we marked them for further interpretation of the classifier performance. To mark germline variants in ICGC samples, we downloaded the corresponding germline VCF files from the ICGC portal. Since germline variants for TCGA-LAML samples were not reported, we called them ourselves with the GATK v.4 pipeline for germline short variant discovery (McKenna *et al*., 2010) using variant quality score recalibration according to GATK Best Practices recommendations (Van der Auwera *et al*., 2013; DePristo *et al*., 2011).

### 2.2 gnomAD pre-filtering

A Mutect2 run on a WGS sample generates about 3.8-4.2 million SNPs and artefacts, including about 3.2-3.4 million germline variants. The number of somatic SNPs is usually far lower (see Table 1). While read qualities and read flags can be used to filter out artefacts, they can hardly help to distinguish somatic and germline variants. Instead, mutational signatures (Figures S1 and S2) and variant allele fraction (VAF) profiles (Figure S3) can be considered. However, since the mutational signatures and VAF distributions for somatic and germline variants overlap, these features can not provide a complete class separation. Hence, classification might be improved by supplying some prior information. For example, it has been proposed (Kalatskaya *et al*., 2017) to mark candidate variants encountered in public databases of germline mutations, such as dbSNP common (Sherry *et al*., 1999) or gnomAD (Karczewski *et al*., 2020). In the present study, we consider pre-filtering of SNP variants using the gnomAD v.3.1.2 database (Karczewski *et al*., 2020). Note that such filtering will partially remove not only germline variants and artefacts but also somatic SNPs (see Table 1). Choosing population allele frequency (AF) cutoff permits tuning the number of variants to exclude.

To determine the optimal gnomAD AF cutoff, we estimated the fraction of SNPs that are retained when removing all variants with gnomAD population AF above a certain threshold. The results for the TCGA-LAML dataset are shown in Figure 1a. As expected, germline variants are reduced in the largest proportion. The maximal gnomAD AF of about 1% might be the optimal threshold when maximizing the somatic recall since at this cutoff the majority of germline variants is already removed whereas somatic variants are almost not affected.

**Fig. 1.**
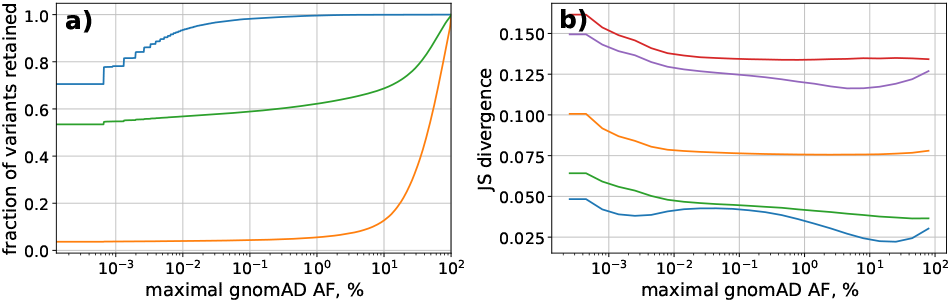
(a) The fraction of somatic (blue), germline (orange) SNPs and artefacts (green) retained in the TCGA-LAML dataset for different gnomAD AF cutoffs. The most significant reduction is observed for germline variants. (b) JS divergence between somatic and germline mutational signatures at different gnomAD AF cutoffs for TCGA-LAML (blue), GACA-CN (orange), LINC-JP (green), ESAD-UK (red), BLCA-US (purple). For all datasets, increasing the AF cutoff reduces the JS divergence, i.e. leads to closer mutational signatures.

It is also important to understand how gnomAD filtering affects mutational signatures. Figure 1b shows Jensen-Shannon (JS) divergence between somatic and germline mutational signatures for different gnomAD AF cutoffs. It can be clearly seen that for all the datasets the JS divergence decreases with the increase of the gnomAD AF cutoff, i.e. the signatures get closer to each other when fewer variants are filtered out. Therefore, one would expect a worse classifier performance for higher AF cutoffs.

So, higher gnomAD AF cutoffs will not only lead to a greater class imbalance but might also worsen the classifier performance. Since somatic variants present in gnomAD are likely to be harmless passenger mutations, we decided to exclude all variants which have a gnomAD record.

### 2.3 Train/test split

For each dataset, we allocated 80% of WGS samples to the train set and 20% of WGS samples to the test set and split the variants accordingly. The neural network classifier was first trained with 70K SNPs of the negative class and 70K SNPs of the positive class randomly sampled from the train set. The classifier was then evaluated on 70K SNPs of the negative class and, where possible, 70K SNPs of the positive class randomly sampled from the test set.

The initial number of somatic SNPs in the TCGA-LAML dataset was below 70K. To achieve the desired number of train instances, data augmentation was applied to the positive class during training.

Our approach to data augmentation is based on the assumption that for a candidate variant of a given origin (e.g. somatic/germline/artefact) such features as read qualities, read flags and VAF are independent of the genomic position of the candidate variant and the corresponding reference sequence. Throughout data augmentation, a new somatic variant is constructed by picking read base qualities, read flags and VAF information from one randomly chosen somatic variant and the mutational context from another one.

### 2.4 Variant encoding

Before passing to the neural network classifier, SNP variants were encoded following the approach in (Friedman *et al*., 2020). First, reads covering the variant site were collected. Unmapped reads, reads that fail platform/vendor quality checks, PCR or optical duplicates were removed. Each read was then converted to a matrix of size *C* × *L*, where *C*=14 is the number of channels encoding the reference sequence and read features, *L* is the read length. The *C*=14 channels were composed of: 4 channels for one-hot encoding of the reference sequence, 4 channels for p-hot encoding of read base qualities (Friedman *et al*., 2020), 6 channels for encoding of read flags (same for all positions in a given read). The following read flags were considered: read mapped in proper pair (0×2), mate unmapped (0×8), read reverse strand (0×10), mate reverse strand (0×20), not primary alignment (0×100), supplementary alignment (0×800). Afterwards, the reads were piled up at the variant site and stacked as a tensor with dimensions *C* × *W* × *DP*, where *DP* is the read depth at the variant site, *W* =150 is the length of the sequence region of interest (ROI) around the variant, the variant column being placed in the center of ROI. While the read depth *DP* may not be the same at different variant sites, the neural network input should have fixed dimensions. So, we imposed the tensor height *H*=70 (around the 80th percentile of read depth distribution for most datasets). Tensors with *DP < H* were padded to the full height *H* using dummy reads with all channels filled with zeros. For tensors with *DP* >*H*, the reads were first sorted by position and then the tensor was cropped by removing reads from the top and the bottom of the stack. Finally, all reads in the tensor were sorted by the base in the variant column (Friedman *et al*., 2020).

### 2.5 Classifier model

The model architecture is shown in Figure 2. The neural network input consists of a variant tensor and variant meta information. The variant tensor is fed into a convolutional block which includes 4 convolutional layers. The output of the convolutional block is flattened out and concatenated with the meta information. The concatenation is followed by 3 dense layers. The output of the sigmoid activation function in the final layer can be used to determine whether a given variant is somatic.

**Fig. 2.**
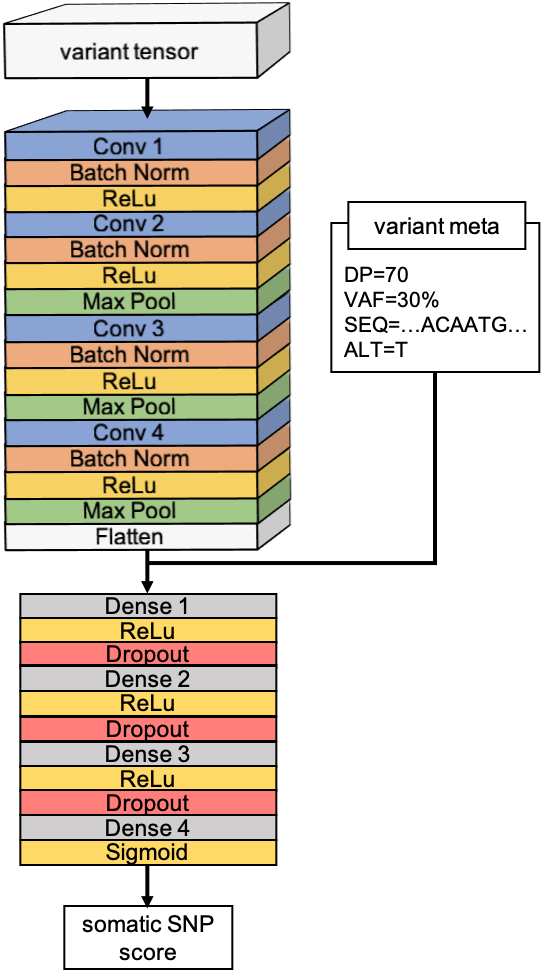
Neural network classifier. Each variant is represented by a variant tensor and variant meta information, the latter being added explicitly to the output of the convolutional block. The output of the sigmoid activation function in the final layer can be used to determine whether a given variant is somatic.

The variant meta information includes the variant allele fraction (VAF), the read depth (DP) and the mutational context. The mutational context is defined as a part of the reference sequence of length *W* around the variant site plus the alternative allele.

For each candidate variant, the VAF is encoded as a fraction of reads with the variant allele. The DP is encoded as the ratio between the number of reads in the variant and the predefined tensor height *H*. When the variant tensor is to be cropped, the VAF and DP are computed before cropping. For the mutational context, one-hot encoding is used.

### 2.6 Performance metric

As the performance metric, we chose the area under receiver operating characteristic curve (ROC AUC). The ROC AUC score does not depend on the class ratio in the dataset and characterize the intrinsic ability of the classifier to separate the two classes. Based on the ROC AUC, classifier performance on datasets with different class ratios can be easily compared.

### 2.7 Neural Network training

The model was implemented with the PyTorch framework (Paszke *et al*., 2019). Optimization of model parameters was assured by the AdamW algorithm (Loshchilov and Hutter, 2018). To speed up the transfer of training data over our computational cluster, we first grouped variant tensors of the same class in sets of 4. At each training iteration, mini-batches of 8 sets were presented to the neural network and the gradient of the binary cross-entropy loss function with respect to model parameters was computed. The model parameters were then updated according to AdamW rules, based on the computed gradient, predefined learning rate and weight decay. To reduce the training time, we used PyTorch automatic mixed precision training.

Hyper-parameter tuning was performed to maximize the classifier performance measured as the ROC AUC. This procedure was performed via a random search (Bergstra and Bengio, 2012), separately for each dataset. During hyper-parameter tuning, the model was trained on 80% of the initial train set and the ROC AUC score was computed on the remaining 20%. The search range of hyper-parameters and their final values are summarized in Table 2. We also attempted to improve the performance by replacing the standard binary cross entropy loss function with the focal loss (Lin *et al*., 2017), but this resulted in worse ROC AUC scores. The final values of hyper-parameters turned out to be the same for all datasets.

**Table 2.** Search range and final values of model hyper-parameters. The final values of hyper-parameters are the same for all datasets.

| Hyper-parameter | Search range | Final value |
| --- | --- | --- |
| Mini-batch size | [4,8,16,32,64] | 8 |
| Learning rate | 1e-8...1e-2 | 1e-3 |
| Weight decay | 1e-6...1e-1 | 1e-1 |
| Dropout | 0.1...0.7 | 0.5 |
| Convolutional kernel size | [3,5] | 3 |
| Num. of conv. channels | [(16,16,16,16), (32,16,32,16),...] | (32,32,32,32) |
| Num. of units in dense layers | [(256,128,64), (128,64,64), ...] | (256,256,128) |

For all datasets, the ROC AUC stabilized after the first 15 epochs when training with the final hyper-parameters. The initial learning rate was then reduced by a factor of 10 and the training continued for 5 more epochs until convergence.

For each dataset, the model was retrained on the train set, then the ROC AUC score was reported on the test set. The average training time was about 19 hours on NVIDIA Tesla V100S.

### 2.8 Mutational signatures

Mutational signatures of SNP variants were constructed by computing the probability of each possible mutation defined as the 3bp-window around the variant site plus the alternative allele (Alexandrov *et al*., 2013).

To quantify the difference between two mutational signatures *P* and *Q*, we used the Jensen-Shannon (JS) divergence:

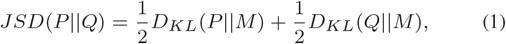

where *M* = 1*/*2(*P* + *Q*) and *D*_*KL*_(*P* ||*Q*) is the Kullback–Leibler divergence:

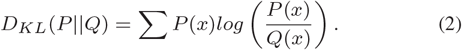

If the opposite is not stated explicitly, the reported JS divergence was computed on mutational signatures obtained after removing variants encountered in gnomAD v.3.1.2 and before normalizing the signatures by trinucleotide reoccurrence in the reference genome.

## 3 Results and Discussion

Neural network performance scores and examples of ROC curves obtained on the independent test set that was not used during training are demonstrated for each dataset in Figure 3a,b. The average ROC AUC test score on all datasets is 0.96±0.01. High ROC AUC scores for all datasets indicate that the proposed neural network classifier is effective for a variety of cancer types and sequencing protocols.

**Fig. 3.**
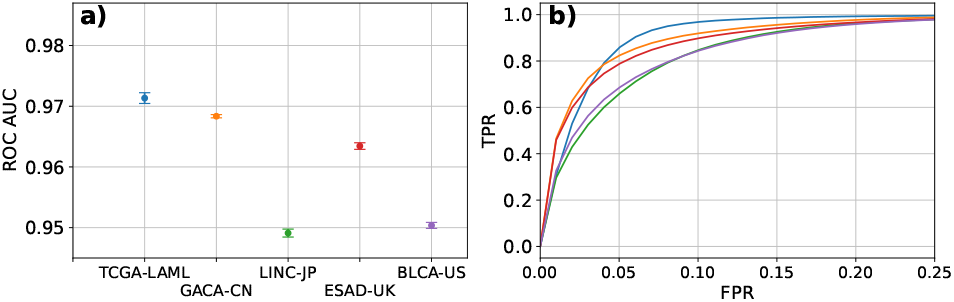
Neural network classifier performance on different datasets. (a) Test ROC AUC scores. (b) Example ROC curves for TCGA-LAML (blue), GACA-CN (orange), LINC-JP (green), ESAD-UK (red), BLCA-US (purple). High ROC AUC values for all datasets indicate that the neural network is able to classify the majority of test SNPs correctly.

Figure 4 shows ROC AUC scores computed separately for somatic vs germline variants classification (a,b) and for somatic variants vs artefacts classification (c,d) as a function of the JS divergence between VAF distributions (a,c) and the JS divergence between mutational signatures (b,d). As can be observed, the classifier is able to filter out germline variants as well as to separate somatic SNPs from artefacts. In general, however, it is more challenging to separate somatic and germline variants than somatic variants and artefacts. It is also easy to note that the JS divergence between VAF distributions is correlated with the eventual neural network performance.

**Fig. 4.**
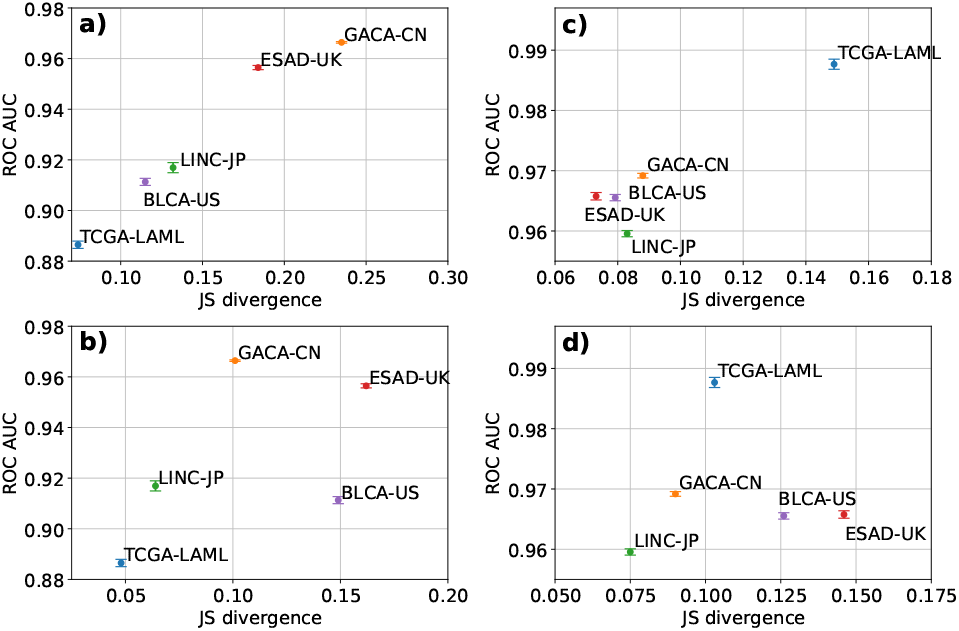
Neural network classifier performance for somatic vs germline variants classification (a,b) and for somatic variants vs artefacts classification (c,d) as a function of the JS divergence between VAF distributions (a,c) and the JS divergence between mutational signatures (b,d). The classifier achieves a better performance when separating somatic variants and artefacts compared to separating somatic and germline variants.

Figure 4 might also indicate that the VAF is more important for classification than the mutational context: although mutational signatures of different kinds of variants are less similar in ESAD-UK and BLCA-US compared to GACA-CN, the neural network scores better on GACA-CN, probably due to the larger divergence between its VAF distributions. On the other hand, the larger difference between mutational signatures in BLCA-US and ESAD-UK compared to LINC-JP leads to a slightly better neural network performance when filtering out artefacts (Figure 4c,d).

Note that in contrast to ISOWN (Kalatskaya *et al*., 2017), the proposed approach not only effectively removes germline variants but also filters out artefacts. In addition, the neural network classifier proves to be effective on samples whose average coverage is much lower (Table 1) than required by VAF-only based tools, such as as SomVarIUS (Smith *et al*., 2016).

In Section 2.2 it was supposed that high gnomAD population AF cutoffs might result in a degraded classifier performance. To verify this assumption, we trained and evaluated the neural network classifier on the TCGA-LAML dataset for two extra gnomAD AF cutoffs: max. gnomAD AF=0.05% and max. gnomAD AF=10%. The resulting ROC AUC scores for somatic vs germline classification (0.853±0.001 and 0.741±0.006 correspondingly) reveal an inferior neural network performance compared to training when all gnomAD variants are excluded (ROC AUC=0.889±0.001, Figure 4a,b). So, raising the gnomAD population AF cutoff indeed leads to a suboptimal performance, presumably due to a stronger similarity between somatic and germline mutational signatures (Figure 1b).

As noted in Section 2.3, we used data augmentation to compensate for the small number of somatic SNPs in the original TCGA-LAML dataset. To check whether generating new examples through data augmentation is necessary to improve the neural network performance, we also trained the classifier by simply equilibrating the classes in the original dataset through upsampling. This resulted in the same performance as on the augmented dataset. Hence, data augmentation did not provide any new information relevant for classification, either because the variants generated through augmentation were too correlated with original SNPs or because all the relevant information was already included in the original dataset.

As noted in the Introduction, the mutational context might provide information relevant for variant classification. As far as SNP variants are concerned, the mutational context is defined with the reference sequence ROI around the variant position plus the alternative allele. However, the role of the ROI length *W* has not been studied yet.

Information about the mutational context is available to the neural network classifier through the variant tensor as well as through the meta information channel connected to the output of the last convolutional layer (Figure 2).

To look into the effect of *W*, we first repeated our experiments without adding the variant meta information explicitly after the convolutional block. This simplified architecture demonstrated the same performance as the initial model (Figure 3). This indicates that all relevant information, including the mutational context, is already extracted from the variant tensor in the convolutional block and explicit addition of variant meta information is in fact not required. This result is in the agreement with (Friedman *et al*., 2020), where explicit addition of variant annotations did not improve artefacts filtering.

Using the simplified architecture, we studied the model performance as a function of *W*. Figure 5b illustrates the results obtained on the BLCA-US dataset. Note that the model performance for *W* < 24 is not shown in Figure 5b since such small values of *W* can not be tested given the chosen configuration of the convolutional block.

**Fig. 5.**
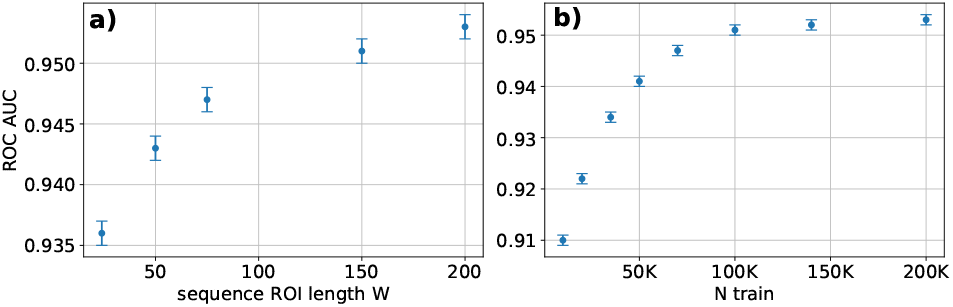
(a) Classifier performance as a function of the sequence ROI length *W* for the BLCA-US dataset (*Ntrain*=70K, *H*=70). Models trained for larger *W* result in a better performance. (b) Classifier performance as a function of the number of train SNP variants *Ntrain* for the BLCA-US dataset (*W* =200, *H*=70). Models trained on more variants result in a better performance.

As shown in Figure 5b, the ROC AUC score gradually improves with *W*. The observed performance improvement might be due to a better artefacts filtering as larger *W* allow for a more accurate estimation of the quality of each read. Note that SNP variants has a 1-bp length, and the effect of *W* could be more pronounced if INDEL variants of different lengths were also considered.

To get a deeper insight on the effect of *W*, we generated neural network saliency maps (Simonyan *et al*., 2013) for several SNP variants (Figure S4). These saliency maps indicate that most of the information relevant for classification is represented by the 3-5bp window around the SNP position.

In Section 2.2 it was noted that complete class separation based on VAF and mutational context is not possible. In this regard, it might expected that classification could be improved by including features that can not be inferred from the variant tensor. For example, the ISOWN method considers variant effect annotation when classifying germline and somatic variants (Kalatskaya *et al*., 2017). To check whether variant effect annotation can improve the model performance, we annotated the SNP variants using the snpEff toolbox (Cingolani *et al*., 2012). The one-hot encoded annotations were then added to the output of the convolutional block. The resulting classifier did not demonstrate any performance improvement.

As noted in Section 2.3, quite a large number of SNP variants was used to train the neural network. When the training time should be reduced or when such a large number of SNPs is unavailable, training on smaller datasets can be considered. However, reducing the train set size often degrades the performance of a machine learning model. Figure 5b shows ROC AUC scores obtained on the BLCA-US dataset for different number of train SNPs *N*_*train*_.

It can be clearly seen that the model performance improves very slowly for *N*_*train*_ > 100K. To understand why the performance is degraded for smaller *N*_*train*_, note that when all variants present in gnomAD are removed, the ratio between germline variants and artefacts becomes about 1:5. When only 5K SNPs of the negative class are used for training (*N*_*train*_ = 10K), the number of germline variants might be too low to adequately represent the mutational signature and the VAF profile.

To assess the biological significance of the somatic variant calls obtained with the neural network approach, we computed the fraction of somatic and non-somatic (germline variants plus artefacts) SNPs in cancer gene census (CGC) genes (Sondka *et al*., 2018) before and after classification. For this analysis, we chose only SNPs labelled as high-impact mutations by the snpEff variant annotation toolbox (Cingolani *et al*., 2012). As usual, we also removed all SNPs catalogued in the gnomAD v.3.1.2 database (Karczewski *et al*., 2020). As a baseline, the percentage of SNPs in CGC genes computed on true labels (before classification) is shown in Table S1 for all datasets. In general, it is significantly greater for somatic variants than for non-somatic SNPs. Since the proportion of non-somatic variants (SNPs) in CGC genes should not depend on the underlying cancer type, we consider the average result (4.9±0.5%) as a baseline for non-somatic SNPs. Note that when computing the overall percentage of SNPs (somatic and non-somatic) in CGC genes per WGS sample, the resulting proportion is closer to that of non-somatic SNPs because even after applying gnomAD and snpEff filtering the number of non-somatic SNPs per WGS sample exceeds the number of somatic SNPs.

The corresponding proportions after classification are shown in Figure 6. For classification, we used the threshold of 0.6 which provides TPR≈90% and FPR≈90% for all datasets. Although the CGC fraction of SNPs classified as somatic is lower compared to the ground truth (Table S1), it is significantly above the designated baseline for non-somatic variants (4.9±0.5%). This provides additional evidence that the neural network can successfully filter out germline variants and artefacts and identify somatic variants in genes that are relevant for cancer biology. For all datasets except ESAD-UK, where this is also not the case for the ground truth proportions, the CGC fraction of SNPs classified as somatic also exceeds the CGC fraction of SNPs classified as non-somatic.

**Fig. 6.**
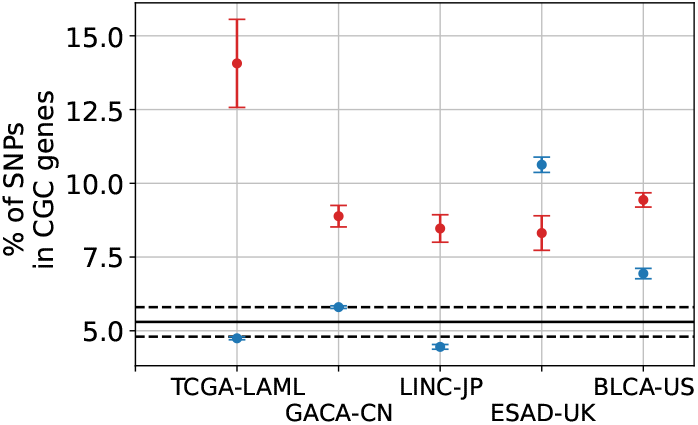
Proportions in CGC genes for SNPs classified as somatic (red) and non-somatic (blue) by the neural network (classification threshold 0.6, with TPR≈90% and FPR≈90%). The baseline for non-somatic variants and its 95% confidence interval are shown with the solid black and dashed lines correspondingly. The predominating proportion of somatic SNPs in CGC genes provides additional evidence that the neural network can successfully filter out germline variants and artefacts.

## 4 Conclusion

In this work, we presented a neural network-based approach for somatic SNP filtering in WGS samples without a matched normal. The method involves simple data preprocessing, can be applied to samples with a typical WGS coverage of around 30x and does not need any manual tuning of filtering parameters. The method is disease-specific and requires training data which can be obtained with standard variant calling tools on several tumor samples for which a matched normal is available.

Due to the large input class inbalance, the proposed method would still necessitate additional steps to filter out germline SNPs and artefacts identified as somatic variants. Although the number of false positives could be reduced by analysing variant reoccurence in cohorts of tumor samples, this approach is not relevant when looking for rare variants. So, future work should include a search for new features that would improve the performance of the neural network classifier *per se*. The proposed approach can also be extended to detection of somatic INDELs or copy number variations in tumor-only setting.

Our implementation of the presented method is freely available as a GitHub repository.

## Acknowledgements

We would like to acknowledge insightful discussions with Julien Gagneur and Stephan Hutter.

