## Supplementary Materials for "Neural network approach to somatic SNP calling in WGS samples without a matched control"

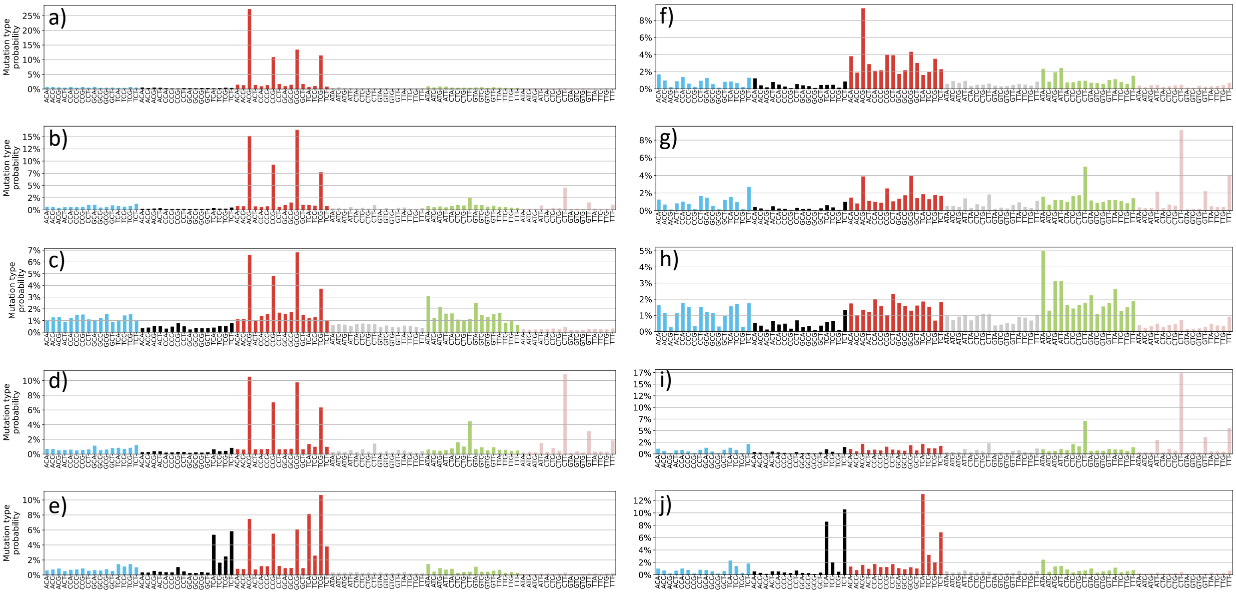


**Figure S1**. Normalized (a-e) and non-normalized (f-j) mutational signatures of somatic variants in the TCGA-LAML (a, f), GACA-CN (b, g), LINC-JP (c, h), ESAD-UK (d, i), BLCA-US (e, j) datasets.

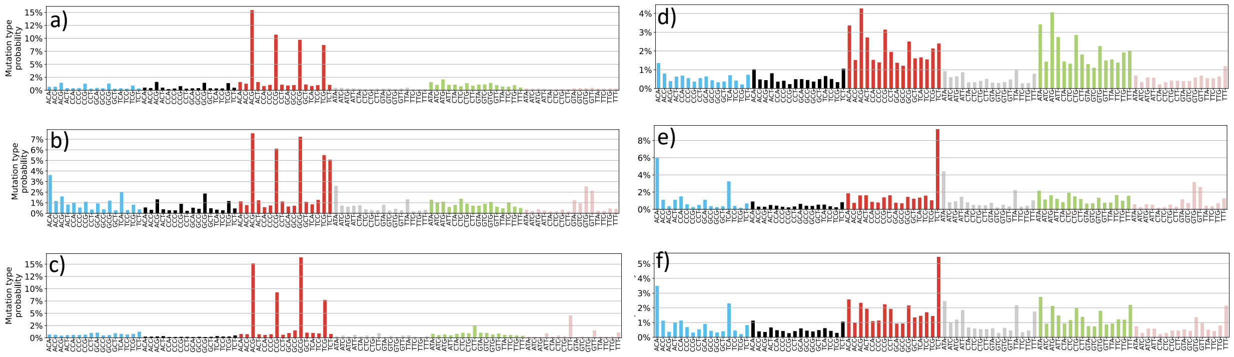

**Figure S2**. Normalized (a-c) and non-normalized (d-f) mutational signatures of germline variants (a, d), sequencing artefacts in the TCGA-LAML dataset (b, e), and sequencing artefacts in the ICGC datasets (c, f). Different sequencing equipment result in slightly different artefacts signatures.

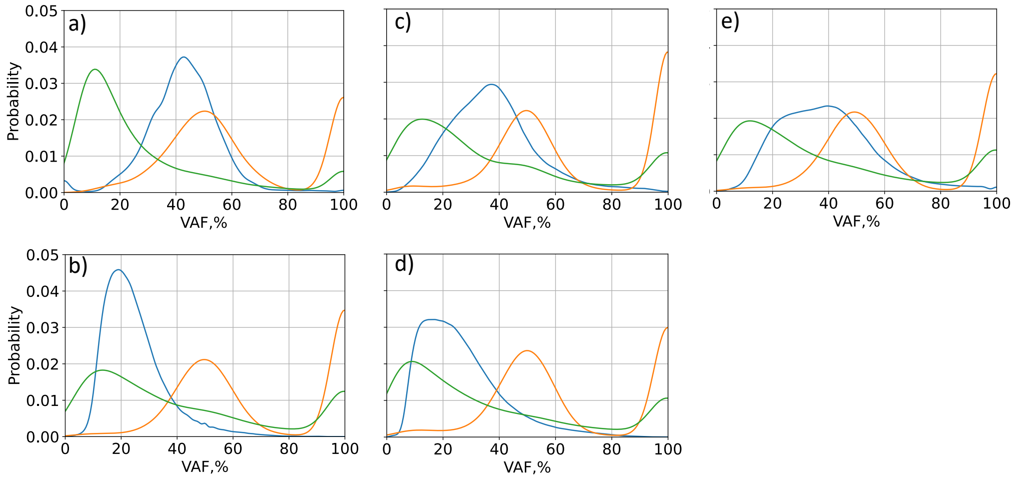


**Figure S3**. Variant allele fraction (VAF) distributions of sequencing artefacts (green), germline variants (red) and somatic variants (blue) in the TCGA-LAML (a), GACA-CN (b), LINC-JP (c), ESAD-UK (d), and BLCA-US (e) datasets.

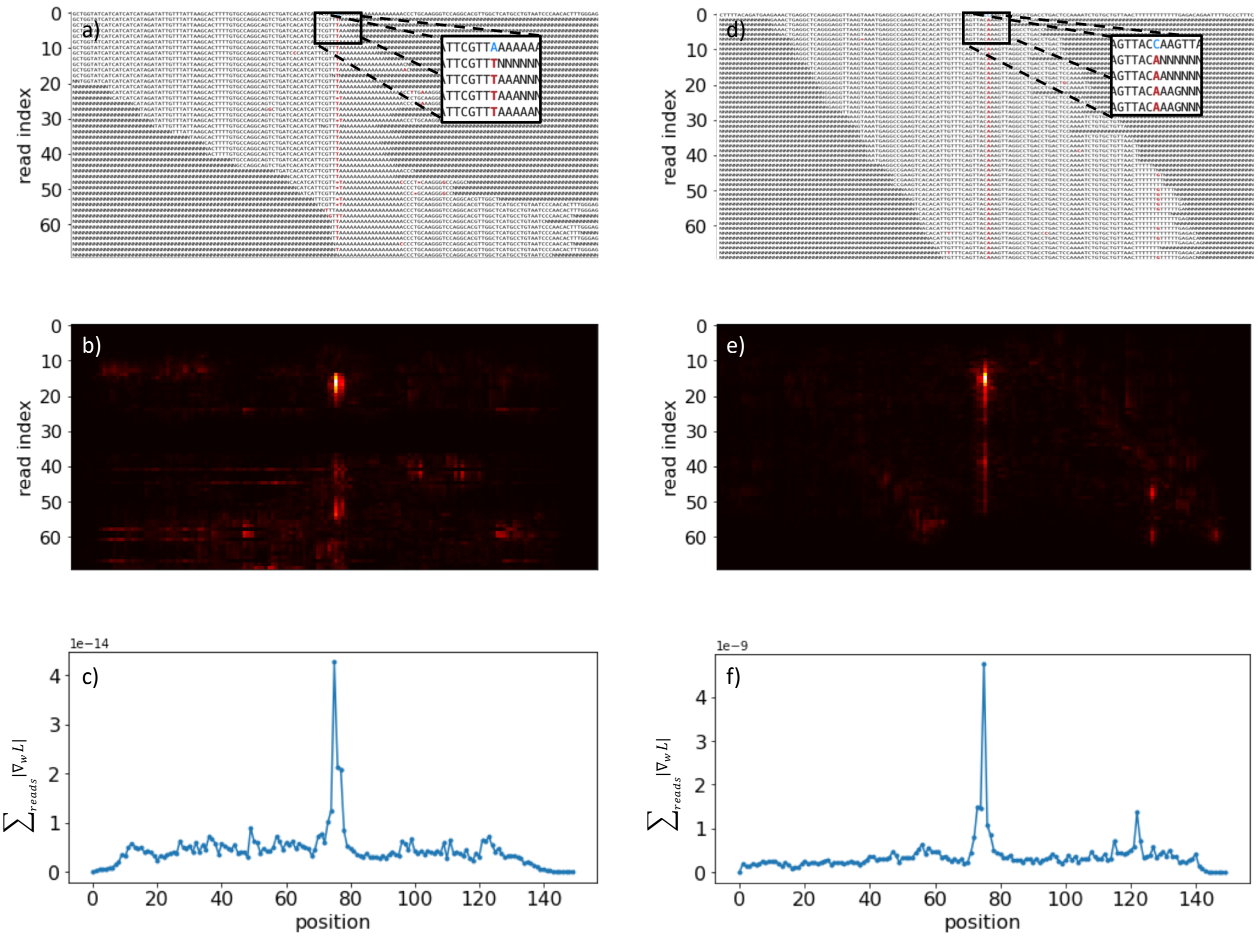


**Figure S4**. Saliency maps for two SNP variants from the BLCA-US dataset. (a,d) –pileup images, (b,e) – saliency maps, brighter regions corresponding to higher gradient amplitudes, (d,f) – sum of (b,e) over the reads axis. Larger gradient amplitudes correspond to a 3-5bp window around the variant site (center of the image).

| Dataset | SOMATIC | | | | NON-SOMATIC | | | | Fisher's test p-value | SNPs in CGC genes (per WGS sample), % |
|---|---|---|---|---|---|---|---|---|---|---|
| | Nsamples | SNPs | SNPs in CGC genes | SNPs in CGC genes (95 % CI), % | Nsamples | SNPs | SNPs in CGC genes | SNPs in CGC genes (95 % CI), % | | |
| BLCA-US | 23 | 484 | 58 | 12±2,9 | 5 | 844 | 51 | 6±1,6 | <0.01 | 6,7 |
| LINC-JP | 28 | 165 | 14 | 8,5±4,3 | 6 | 1692 | 84 | 5±1 | 0.066 | 5,1 |
| ESAD-UK | 41 | 369 | 39 | 10,6±3,1 | 8 | 667 | 66 | 9,9±2,3 | 0.74 | 10 |
| GACA-CN | 25 | 78 | 13 | 16,7±8,3 | 6 | 765 | 41 | 5,4±1,6 | <0.01 | 5,7 |
| TCGA-LAML | 47 | 50 | 13 | 26±12,2 | 10 | 4399 | 202 | 4,6±0,6 | <0.01 | 4,6 |
| Total* | 164 | 1096 | 124 | 11,3±1,9 | 35 | 7700 | 378 | 4,9±0,5 | <0.01 | 5,1 |

*results on TCGA-LAML(somatic) excluded;
results on ESAD-UK (non-somatic) excluded

**Table S1**. Number of somatic and non-somatic (germline+artefacts) SNP variants in each dataset (true labels) after excluding all gnomAD variants and selecting only SNPs associated with a "HIGH" impact (according to snpEff annotations). For each variant type, absolute number and the relative proportion (with 95% confidence intervals) of SNPs in cancer gene census (CGC) genes are shown. The Fisher's test p-value corresponds to the evidence that the proportion of somatic SNPs located in CGC genes is higher than the proportion of non-somatic SNPs. The last column corresponds to the proportion of SNPs in CGC genes per WGS sample. This proportion is shifted to that of non-somatic variants since the number of non-somatic SNPs outweigh the number of somatic SNPs per WGS sample.
